# Crosstalk Between One Carbon Metabolism and Eph Signaling Promotes Neural Stem Cells Differentiation Through Epigenetic Remodeling

**DOI:** 10.1101/240895

**Authors:** Mohamad-Ali Fawal, Thomas Jungas, Anthony Kischel, Christophe Audouard, Jason S. Iacovoni, Alice Davy

**Affiliations:** Centre de Biologie du Développement (CBD), Centre de Biologie Intégrative (CBI), Université de Toulouse, CNRS, UPS, 118 route de Narbonne, 31062 Toulouse, France.; Bioinformatic Plateau I2MC, INSERM and University of Toulouse, Toulouse, France.

**Keywords:** 1 carbon metabolism, neural progenitors, folate, ephrin, histone methylation, differentiation, stem cells

## Abstract

Metabolic pathways, once seen as a mere consequence of cell states, have emerged as active players in dictating different cellular events such as proliferation, self-renewal and differentiation. Several studies have reported a role for folate-dependent 1-carbon (1C) metabolism in stem cells, however, its exact mode of action and how it interacts with other cues is largely unknown. Here, we report a link between the Eph:ephrin cell-cell communication pathway and 1C metabolism in controlling differentiation of neural stem cells. Transcriptional and functional analyses following ephrin stimulation revealed alterations in folate metabolism-related genes and enzymatic activity. In vitro and in vivo data indicate that Eph-B forward signaling alters the methylation state of H3K4 by regulating 1C metabolism, and locks neural stem cells in a differentiation-ready state. The functional link between cell-cell communication, metabolism and epigenetic remodeling identifies a novel triad in the control of stem cell self-renewal vs. differentiation.

**HIGHLIGHTS:**

- 1C folate metabolism is regulated by local cell-to-cell communication
- Description of Eph-B transcriptional response in NSC
- Eph activation decreases the expression and activity of DHFR
- Inhibition of DHFR modifies epigenetic marks and impairs self-renewal of neural stem cells
- Decreased H3K4 methylation locks neural stem cells in a pro-differentiation state

**eTOC BLURB:** Fawal et al. present evidence that Eph-B forward signaling inhibits 1C folate metabolism in neural stem cells leading to alteration of H3K4 methylation on key progenitor genes. In addition, they show that these epigenetic changes are inherited and maintained in the long term, thus locking NSC into a differentiation ready state.

## INTRODUCTION

A distinctive feature of stem cells (SCs) is their dual capacity for self-renewal and multipotent differentiation. Recent evidence has revealed that in addition to growth factors and the extracellular matrix, various metabolic pathways contribute to the balance between self-renewal and differentiation of SCs. Indeed, while defined metabolic states are associated with stemness, a switch in metabolic pathways supports divergent cell fate through coordination with signaling and genetic/epigenetic regulation (Folmes et al., 2012b; Shyh-Chang et al., 2013a; Zhang et al., 2012). In the developing brain, a tight balance between self-renewal and differentiation of neural stem cells (NSC) is important to ensure that correct numbers of neural cells are generated (Götz et al., 2016; Urbán and Guillemot, 2014). While several studies have highlighted an important role for glycolysis, lipogenesis and mitochondrial activity in neurogenesis (Knobloch and Jessberger, 2017), the one-carbon (1C) pathway has comparatively received less attention.

1C metabolism is a universal metabolic pathway that couples purine synthesis (required for DNA replication and cell proliferation) and reactive methyl carrier S-adenosylmethionine (SAM) synthesis which is required for methylation reactions and epigenetic modifications. It is fueled by nutrients present in the serum and in the cerebrospinal fluid and it requires folate as an essential co-factor (Ducker and Rabinowitz, 2017). Folate is best known as a vitamin that prevents neural tube defects during fetal development (Schorah et al., 1980), and whose deficiency contributes to severe neurodevelopmental and neurological disorders such as epilepsy, microcephaly and intellectual disability (Serrano et al., 2012), as well as neurodegenerative diseases such as dementia and Alzheimer’s disease (Malouf et al., 2003; Ramos et al., 2005). Previous studies have highlighted the role of folate in NSC self-renewal and differentiation. For instance, folate deficiency inhibits the proliferation of adult hippocampal NSC in vivo and induces NSC apoptosis in vitro (Kruman et al., 2005; Zhang et al., 2008). Conversely, NSC respond to folate with increased proliferation and neuronal differentiation (Liu et al., 2013; Luo et al., 2013). These reports point out the role of 1C metabolism in NSC regulation, however, whether and how it interacts with other regulatory mechanisms that control neurogenesis remain unknown.

Here, we report a link between the Eph:ephrin cell-cell communication pathway and 1C metabolism. Eph receptors, the largest family of receptor tyrosine kinases and their ligands, the ephrins, enable contact-mediated signaling and participate in a wide spectrum of developmental processes through control of cell migration, adhesion and repulsion. While the role of Eph:ephrin signaling in both embryonic and adults neurogenesis is well documented in vivo (Laussu et al., 2014), its biological outcome is divergent in different contexts (Ashton et al., 2012; Liu et al., 2017; Ottone et al., 2014). In addition, even though downstream effectors of Eph receptors, such as small GTPases, cytoplasmic kinases and phosphatases are well studied in the context of tissue morphogenesis or axon guidance (Kania and Klein, 2016), the molecular mechanisms underlying their effect in neurogenesis remain poorly understood.

To gain further insights, we used in vitro and in vivo analyses of embryonic NSC to identify the molecular mechanisms regulating proliferation, self-renewal and differentiation downstream of Eph. We show that ephrin stimulation impaired NSC self-renewal under SC culture conditions and in the developing neocortex. This impairment was associated with increased differentiation but not reduced proliferation rate. Combination of transcriptional and enzymatic analyses revealed a link between Eph-B signaling and 1C folate pathway. This link was further confirmed in vivo. We present evidence that Eph-B forward signaling through alterations in 1C metabolism modifies H3K4 methylation on key progenitor genes. Finally, we show that these epigenetic changes are inherited and maintained in the long term, thus locking NSC into a differentiation ready state.

## RESULTS

### Eph-B forward signaling regulates DHFR

To identify the downstream effectors of Eph-B forward signaling, we performed transcriptional analyses of NSC after 2 and 6 hours (h) stimulation with ephrin-B1 recombinant proteins (eB1-Fc) which led to transient phosphorylation of Eph-B receptors (Figure S1A). About 400 genes were differentially expressed upon Eph-B activation (Figure S1B and C) and gene ontology annotation revealed enrichment in genes involved in regulation of gene expression, cellular component organization, metabolic and cellular processes (Figure 1A). Upon closer inspection, we identified several genes from the 1C folate pathway whose expression was decreased (Figure S1D) suggesting that this pathway may be downstream of Eph-B signaling. Some of these target genes were validated by qRT-PCR and among them, *Dhfr* (Dihydrofolate Reductase, DHFR) was consistently and significantly downregulated following eB1-Fc treatment (Figure 1B). This decrease in DHFR mRNA at 6h correlated with a detectable and significant decrease of DHFR protein levels 72h post-eB1-Fc treatment (Figure 1C). DHFR is a key enzyme in the 1C metabolic pathway that catalyzes the reduction of 7,8-dihydrofolate (DHF) to 5,6,7,8-tetrahydrofolate (THF) and whose activity is inhibited by Methotrexate (MTX), an anti-folate compound (Goldin et al., 1955). To evaluate the impact of Eph signaling on DHFR activity, we measured DHFR activity in NSC following eB1-Fc treatment and compared it to MTX treatment. Interestingly, eB1-Fc treatment inhibited DHFR activity in NSC as efficiently as MTX, but with slower kinetics (Figure 1D). The delay in DHFR inhibition following eB1-Fc treatment and the sequential decrease of *Dhfr* mRNA preceding its simultaneous protein and activity reduction indicates that activation of Eph-B forward signaling inhibits DHFR activity by regulation of its expression rather than by acting directly on its activity.

**Figure 1.**
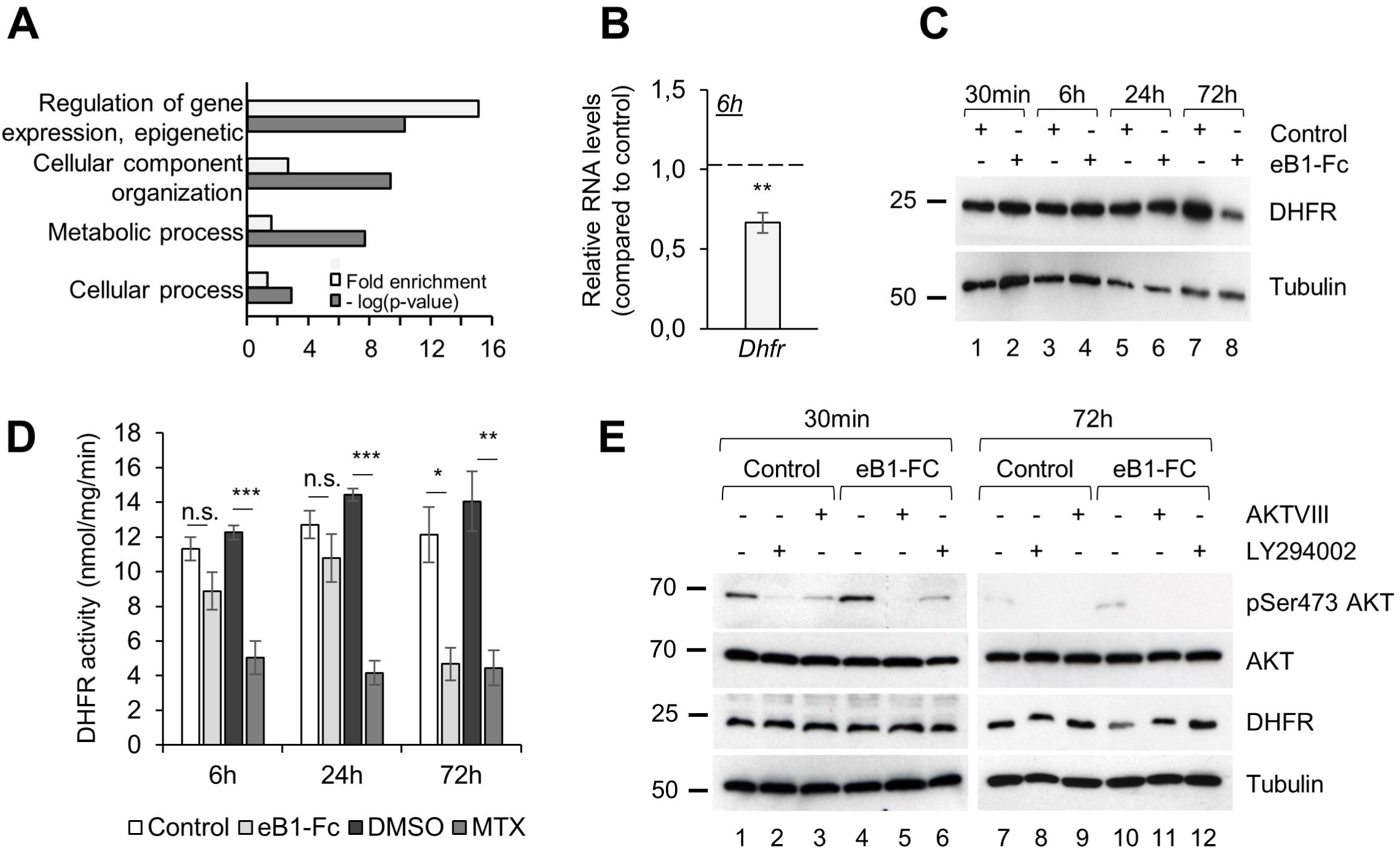
Eph-B forward signaling regulates DHFR. (A) Gene ontology enrichment analysis of differentially expressed genes following eB1-Fc treatment. (B) qRT-PCR of cultured NSC treated for 6h with eB1-Fc. Relative *dhfr* mRNA expression is shown. (C) NSC were treated as in (B) for the indicated time. Samples were analyzed by western blot using the indicated primary antibodies. (D) DHFR activity in NSC lysates treated with either with eB1-Fc or MTX for the indicated times. (E) NSC were treated with either AKT VIII or LY294002 inhibitors for 30min and 72h then analyzed by western blot with the indicated antibodies. Statistical analysis was performed using Mann Whitney test (B) and one-way ANOVA test followed by the Bonferroni method (D). Data are reported as mean ± SEM (*P < 0.05; **P < 0.01; *** P<0;005).

Intracellular effectors downstream of Eph-B have been described in several contexts (Kania and Klein, 2016), however the mechanisms underlying their effect on transcriptional programs have yet to be fully understood. One of the main candidate to fulfill the mediator role is AKT which acts as a hub controlling the activity of a plethora of transcription factors (Manning and Cantley, 2007). Upon eB1-Fc treatment, AKT is phosphorylated with similar kinetics to those observed for Eph-B (Figure S1E) indicating that AKT is an effector of Eph-B forward signaling in NSC. To test this, we incubated NSC with AKT VIII or LY294002, two AKT inhibitors which efficiently inhibited AKT phosphorylation (Figure 1E). Interestingly, AKT inhibition prevented the decrease in DHFR protein levels 72h post-eB1-Fc treatment (Figure 1E). Thus, our data indicate that Eph-B forward signaling leads to a decrease in DHFR protein level via AKT.

### Eph-B Forward signaling impairs NSC self-renewal and promotes their differentiation

To test whether Eph signaling and DHFR inhibition are relevant for NSC maintenance, we tested the capacity of these cells to form secondary spheres which is an indicator of self-renewal. We treated NSC with a single dose of eB1-Fc with or without folinic acid (FNA) which replenishes the THF pools depleted by DHFR inhibition (Ortiz et al., 2000; van Ede et al., 2001). eB1-Fc treatment led to a decreased number of secondary spheres which was rescued by FNA supplementation (Figure 2A). Similarly, inhibition of DHFR by MTX treatment also led to a decrease in the number of secondary spheres which was rescued by FNA (Figure S2A). These results indicate that inhibition of DHFR leads to a decrease in the self-renewing potential of NSC.

**Figure 2.**
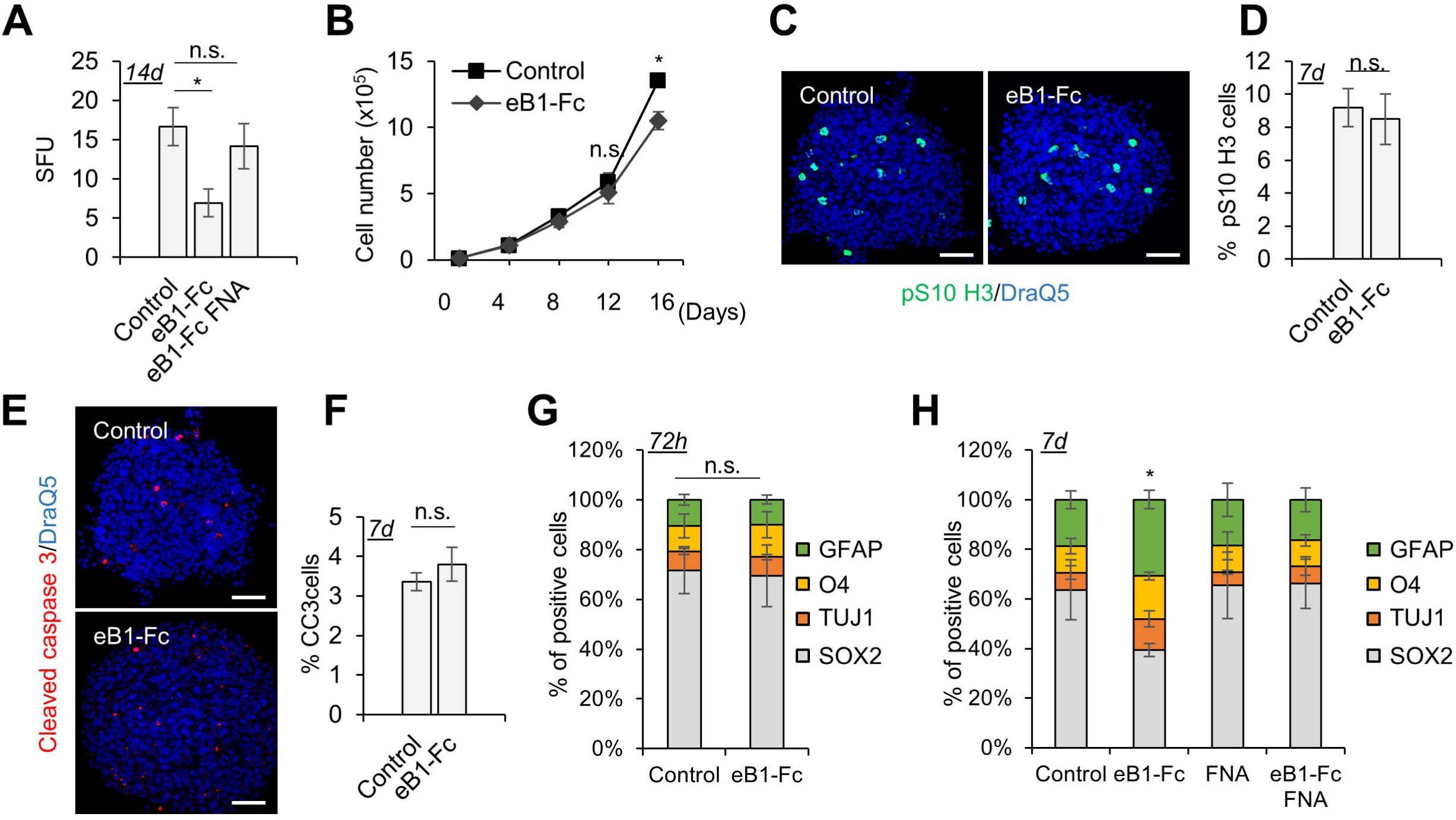
Eph-B forward signaling impairs NSC self-renewal by promoting their differentiation. (A) Neurospheres were dissociated and incubated for 7 days with either eB1-Fc or eB1-Fc supplemented with FNA as indicated. Neurospheres were then dissociated and cultured under stem culture conditions (10^2^ cells/ml) for 1 week and the number of spheres were counted. Sphere-Forming Unit (SFU) is calculated according to the following formula: SFU = (number of spheres counted / number of input cells) * 100. (B) Quantification of cell numbers after 4, 8, 12 and 16d of treatment with eB1-Fc. (C) Representative immunofluoresence images of neurospheres treated with eB1-Fc for 1 week and stained with pS10 H3 (H3P) antibody (green) and DraQ5 (blue). (D) Quantification of H3P positive cells from (C) using Image J. (E) Representative immunofluoresence images of neurospheres treated as in (C) and stained with cleaved caspase 3 (CC3) antibody (red) and DraQ5 (blue). (F) Quantification of CC3 positive cells from (E) using Image J. (G) Quantification by FACS analysis of NSC treated with eB1-Fc for 72h and stained with SOX2, TUJ1, O4 and GFAP antibodies. Statistical significance was performed on SOX2 population. (H) Quantification by FACS analysis of NSC treated with eB1-Fc, FNA alone or eB1-Fc/FNA for 7d and stained with SOX2, TUJ1, O4 and GFAP antibodies. Statistical significance was performed on SOX2 population. h: hours, d: days. Scale bars represent 100 μm. Statistical analysis was performed using one-way ANOVA test followed by the Bonferroni method (A),or Fisher’s LSD test (G,H), multiple t-test using the holm-sidak method (B) and Mann Whitney test (D,F). Data are reported as mean ± SEM (*P < 0.05).

This decrease in self-renewing potential could be due to a decrease in the rate of proliferation or apoptosis. Thus, we tested whether these two parameters were altered following eB1-Fc treatment. Activation of Eph-B forward signaling did not lead to a significant modification in cell number until 16 days (d) post-treatment (Figure 2B) and it had no effect on the mitotic index or apoptosis at 7d (Figure 2C-F). These results suggest that decreased proliferation or increased apoptosis are not the driving factors leading to the change in self-renewing potential observed upon eB1-Fc treatment.

Next, we hypothesized that the decrease in the self-renewing potential induced by Eph-B forward signaling was due to enhanced differentiation. In self-renewing culture conditions neurospheres are heterogeneous due to NSC spontaneous differentiation in neurospheres. In these conditions, we assessed whether activation of Eph signaling is sufficient to promote differentiation. We performed FACS analyses with markers of progenitor cells (SOX2) and differentiated cells (TUJ1, GFAP, O4). While these analyses of eB1-Fc treated NSC showed no significant differences 72h post-treatment (Figure 2G), we observed an expansion of the fraction of differentiated cells at the expense of progenitor cells 7d post-treatment (Figure 2H). In addition, while FNA supplementation alone had no effect on NSC differentiation, it was sufficient to rescue the loss of progenitor cells following eB1-Fc treatment (Figure 2H). Immunofluorescence (IF) analysis confirmed the decrease in progenitor cells (SOX2+) and the increase in neurons (TUJ1+), oligodendrocytes (O4+) and astrocytes (GFAP+) (Figure S2C). Importantly, inhibition of DHFR by MTX resulted in a similar decrease in the progenitor fraction and an increase in the differentiated fraction that could be rescued by FNA supplementation (Figure S2B and D). These results indicate that Eph-B forward signaling decreased the progenitor pool by altering the 1C folate pathway and promoting differentiation at the expense of self-renewal.

### Inhibition of DHFR via Eph-B forward signaling alters epigenetic marks in NSC

One carbon folate pathway has been linked to altered Histone3 methylation (Garcia et al., 2016; Mentch et al., 2015; Shyh-Chang et al., 2013b). We thus analyzed methylation on Histone3, which is known to be required for the maintenance of defined cellular states (Benayoun et al., 2014; Mohamed Ariff et al., 2012), in response to eB1-Fc or MTX treatment as a control. At 72h and prior to any detectable differentiation (Figure 2G), eB1-Fc treatments led to a significant decrease in the H3K4me3 levels and to lesser extents both levels of H3K9me2 and H3K27me3 (Figure 3A). Interestingly, H3K4me3 decrease correlated temporally with the decrease in DHFR activity as no significant change of this mark was observed prior to 72h post-eB1-Fc treatment (Figure S3A). To ascertain that the decrease in H3K4me3 was due to altered 1C folate metabolism, we repeated the experiment in presence of FNA. Supplementation with FNA rescued the decreased H3K4me3 in both eB1-Fc and MTX treatments (Figure 3B). Moreover, inhibition of AKT prevented the decrease in H3K4me3 (Figure S3B) consistent with this mark being downstream of DHFR inhibition.

**Figure 3.**
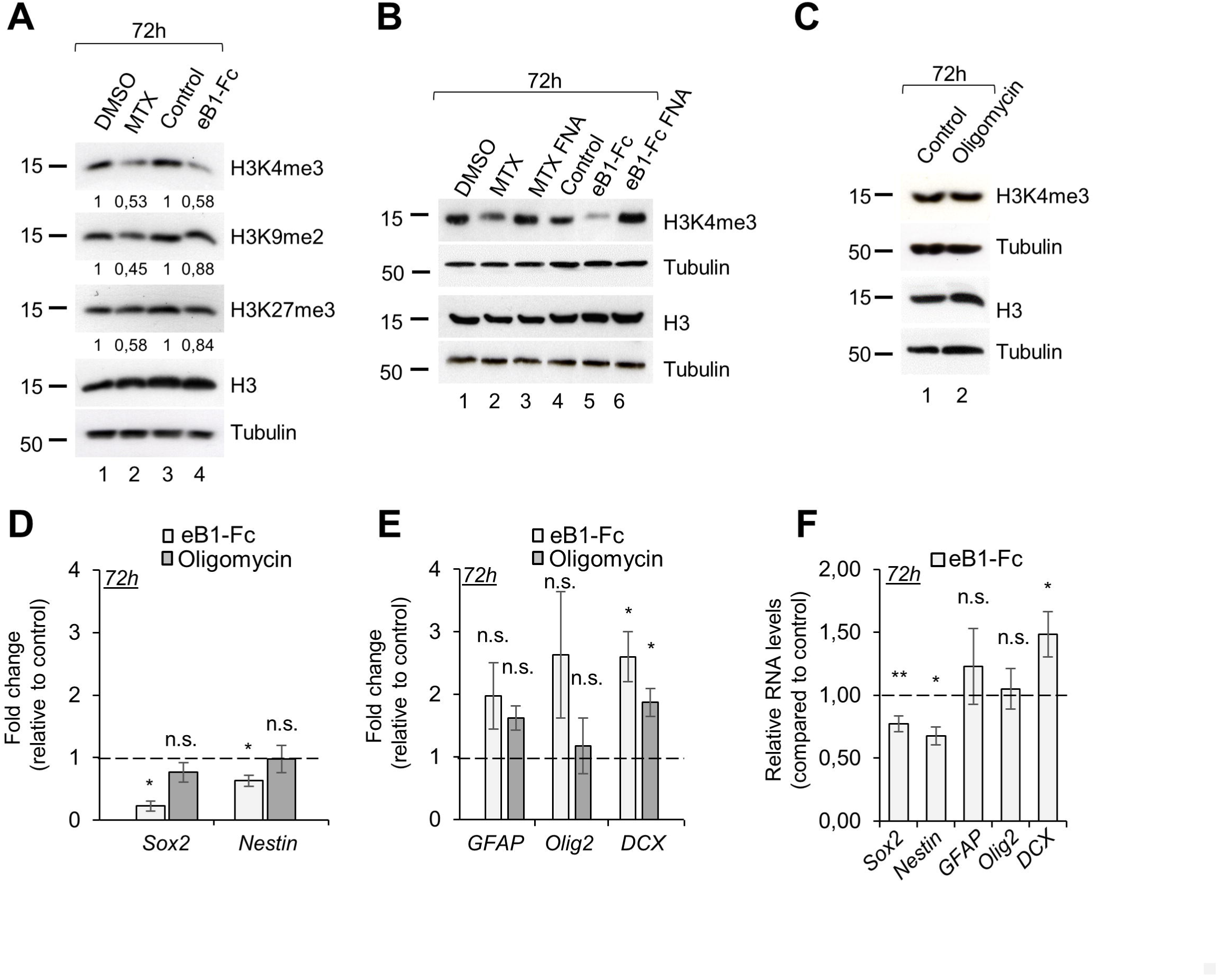
Inhibition of DHFR via Eph-B forward signaling alters epigenetic marks in NSC. (A) Western blot analysis of NSC treated for 72h with MTX or eB1-Fc as indicated. Primary antibodies are indicated on the side. Tubulin and H3 are used as loading controls. (B) Western blot analysis of NSC treated with MTX or eB1-Fc for 72h and supplemented with FNA as indicated. Primary antibodies are indicated on the side. Tubulin and H3 are used as loading controls. (C) Western blot analysis of NSC treated with Oligomycin for 72h. Primary antibodies are indicated on the side. Tubulin and H3 are used as loading controls. (D) NSC were treated with either eB1-Fc or Oligomycin for 72h and H3K4me3 ChIP was performed. The graph shows qRT-PCR analysis of the promoter region of *Sox2* and *Nestin* gene. (E) NSC were treated as in D. The graph shows qRT-PCR analysis of H3K4me3 ChIP of the promoter region of *GFAP*, *Olig2* and *D*cx. (F) qRT-PCR analysis of *Sox2*, *Nestin, GFAP*, *Olig2* and *D*cx mRNA expression 72h following eB1-Fc treatment. Statistical analysis was performed using paired t-test (D, E) and ratio paired t-test (F). Data are reported as mean ± SEM (*P < 0.05; **P < 0.01).

It was recently shown that perturbation of mitochondrial function impacts NSC self-renewal in the developing neocortex (Khacho et al., 2017). In order to assert that the epigenetic changes observed above were the cause and not the consequence of NSC differentiation, we perturbed mitochondrial electron transport chain using Oligomycin, an ATP synthase inhibitor (Liu and Schubert, 2009). As expected, Oligomycin-treatment decreased the NSC capacity to form secondary spheres (Figure S3C). FACS and IF analysis revealed an increase in the differentiated fraction and a decrease in the progenitor fraction following Oligomycin treatment (Figure S3D and E). Importantly, while Oligomycin treatment impaired NSC self-renewal in a manner similar to eB1-Fc treatment, no change in total H3K4me3 levels was detected (Figure 3C). Thus, the decrease in H3K4me3 observed following eB1-Fc treatment is not an indirect consequence of differentiation.

H3K4me3 is important for maintaining euchromatin and is generally associated with promoters of actively transcribed genes (Benayoun et al., 2014). Recent epigenomic profiling of histone methylation patterns in neural progenitors have correlated the presence of H3K4me3 with the expression of crucial transcription factors during neocortex development (Albert et al., 2017). Thus, we performed chromatin immunoprecipitation (ChIP) experiments on NSC to test whether eB1-Fc treatment modify H3K4me3 at specific promoters. Using H3K4me3 ChIP-seq data (Albert et al., 2017; Wu et al., 2011), we designed RT-PCR primers for promoters of progenitor-specific genes (*Sox2* and *Nes* (Nestin)) and differentiation-related genes (*Olig2*, *GFAP*, *Dcx*) and we performed H3K4me3 ChIP following eB1-Fc or Oligomycin treatment. These analyses revealed a decrease of H3K4me3 at the promoters of progenitor specific genes following eB1-Fc treatment but not Oligomycin treatment (Figure 3D). Consistent with our previous data, DHFR inhibition by MTX also led to a decrease in H3K4me3 at promoter regions of progenitor-specific genes (Figure S3F). In contrast, the level of H3K4me3 were either not changed or increased at the promoter of differentiation-related genes in response to eB1-Fc or Oligomycin treatment (Figure 3E), suggesting that this increase was associated with induced differentiation. Importantly, the changes of H3K4me3 at promoters of progenitor and differentiation specific genes in response to eB1-Fc treatment correlated with changes in mRNA levels of these genes (Figure 3F), consistent with the fact that enrichment of H3K4me3 is observed at actively transcribed promoters. These results indicate that DHFR inhibition leads to decreased H3K4 tri-methylation which differentially affects promoters of progenitor vs. differentiation-related genes.

Together, these data show that while Eph-B forward signaling, MTX and Oligomycin treatment lead to a reduction in the progenitor pool, they do so by different mechanisms. This strongly supports the notion that Eph-B forward signaling controls NSC self-renewal through 1C folate mediated modification of epigenetic marks.

### Long term inheritance of the epigenetic marks and differentiation program induced by DHFR inhibition

Changes in histone methylation are generally associated with stable cell fate decisions (Greer and Shi, 2012). To investigate the stability of the epigenetic changes observed upon eB1-Fc treatment, we analyzed H3K4me3 levels at the promoters of progenitor and differentiation-specific genes after dissociation of the treated NSC (first generation) and culture without treatment (second generation) (Figure 4A). Under these conditions, 3 out of 4 selected promoter regions for progenitor-specific genes exhibited decreased H3K4me3 in progeny of cells that had been previously exposed to eB1-Fc treatment (Figure 4B) despite the fact that DHFR activity was no longer inhibited (Figure 4C). As expected, initial Oligomycin treatment, had no effect on the level of H3K4me3 at the promoter of either *Sox2* or *Nes* in NSC progeny (Figure 4B). Similarly, the significant increase in H3K4me3 at the promoter of *Dcx* in response to eB1-Fc treatment was maintained in second generation NSC (Figure 4D). These results indicate that the changes in H3K4me3 at specific promoter regions are inherited from the initial treatment.

**Figure 4.**
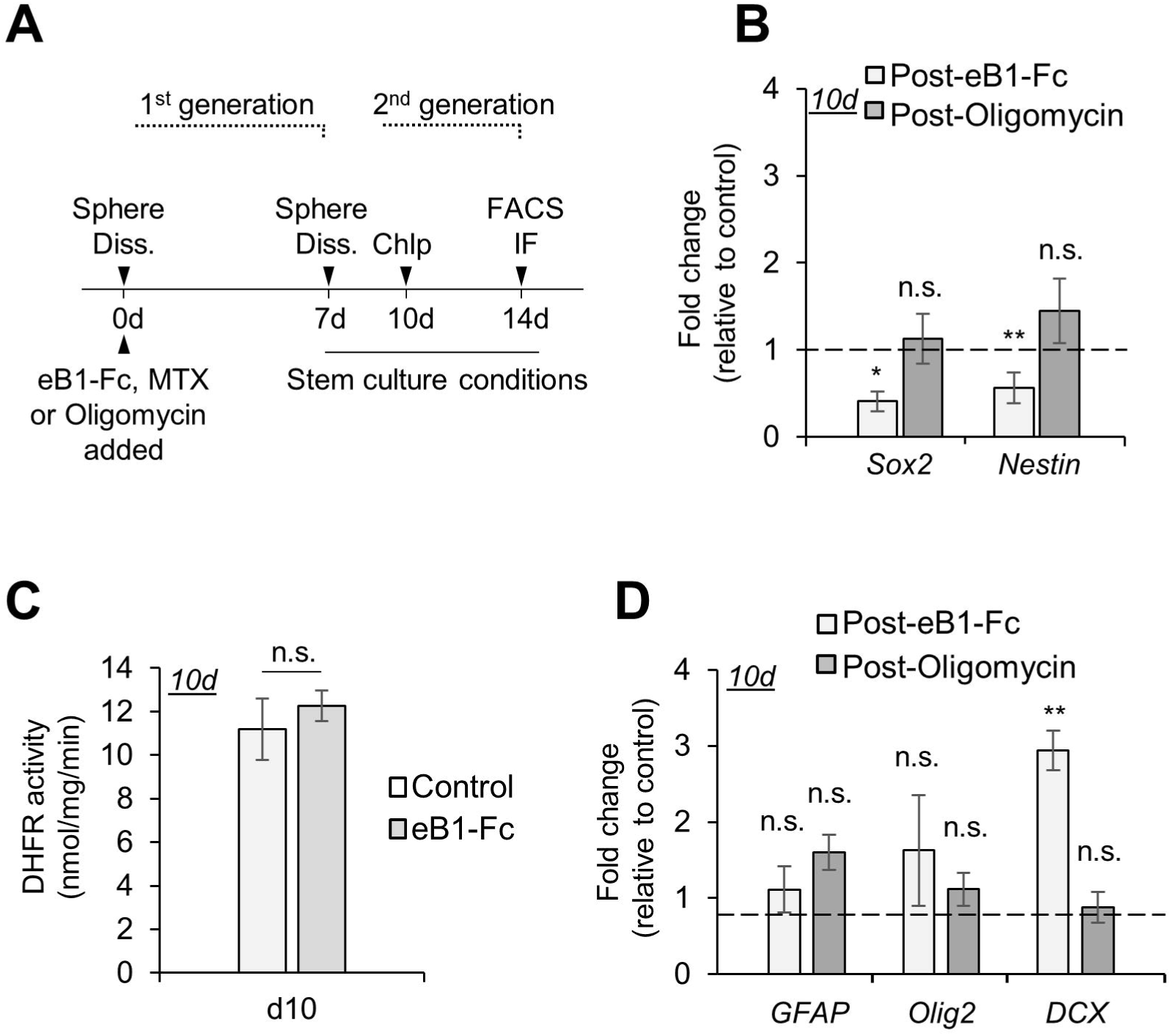
Long term inheritance of the epigenetic marks induced by DHFR inhibition. (A) Graphical presentation of the experimental design used to assess the inheritance of epigenetic marks. NSC were treated and cultured for 7d. Neurospheres were then dissociated and cultured without treatment under stem cell culture conditions and assessed for ChIP, DHFR activity (d10) or FACS, IF (d14). (B) NSC were treated with eB1-Fc or Oligomycin and cultured as in (A). The graph shows the qRT-PCR analysis of the promoter of *Sox2* and *Nestin* following H3K4me3 ChIP. (C) NSC were treated with eB1-Fc or Oligomycin and cultured as in (A). The graph shows the qRT-PCR analysis of the promoter of *GFAP*, *Olig2* and *Dcx* following H3K4me3 ChIP. (D) DHFR activity was measured in NSC treated as in (A). Statistical analysis was performed using paired t-test (B, D) and Mann Whitney test (C). Data are reported as mean ± SEM (*P < 0.05; **P < 0.01).

The inheritance of the changes in H3K4me3 following eB1-Fc initial treatment suggested that the pro-differentiation transcriptional program was also inherited by the progeny. To confirm this hypothesis, we tested the stemness as well as the different cell populations within the second generation neurospheres 14 days post-eB1-Fc treatment. Similar to the first generation, the self-renewal capacity of the secondary NSC was also lower following the initial treatment with eB1-Fc (Figure 5A). This decrease was not a consequence of the overall decrease of progenitors following the initial treatment since Oligomycin treatment that induced a high decrease in the progenitor pool in the first generation (Figure S3D), had no detectable effect on the second generation (Figure 5B). Consistently, this decrease in stemness was also observed by FACS and IF analysis where the decrease in the progenitor fraction correlated with an increase in the fraction of second generation differentiated cells (Figure 5C and S4A) but not with Oligomycin treatment (Figure 5D and S4B). Finally, FNA supplementation of first generation NSC rescued the phenotype observed following eB1-Fc treatment at day 14 (Figure 5C and S4A). These results reveal the co-inheritance of changes in H3K4me3 epigenetic marks and changes in progenitor vs. differentiated fraction following eB1-Fc stimulation. Taken together, these results show that Eph-B forward signaling triggers long term epigenetic changes promoting differentiation and suppressing self-renewal.

**Figure 5.**
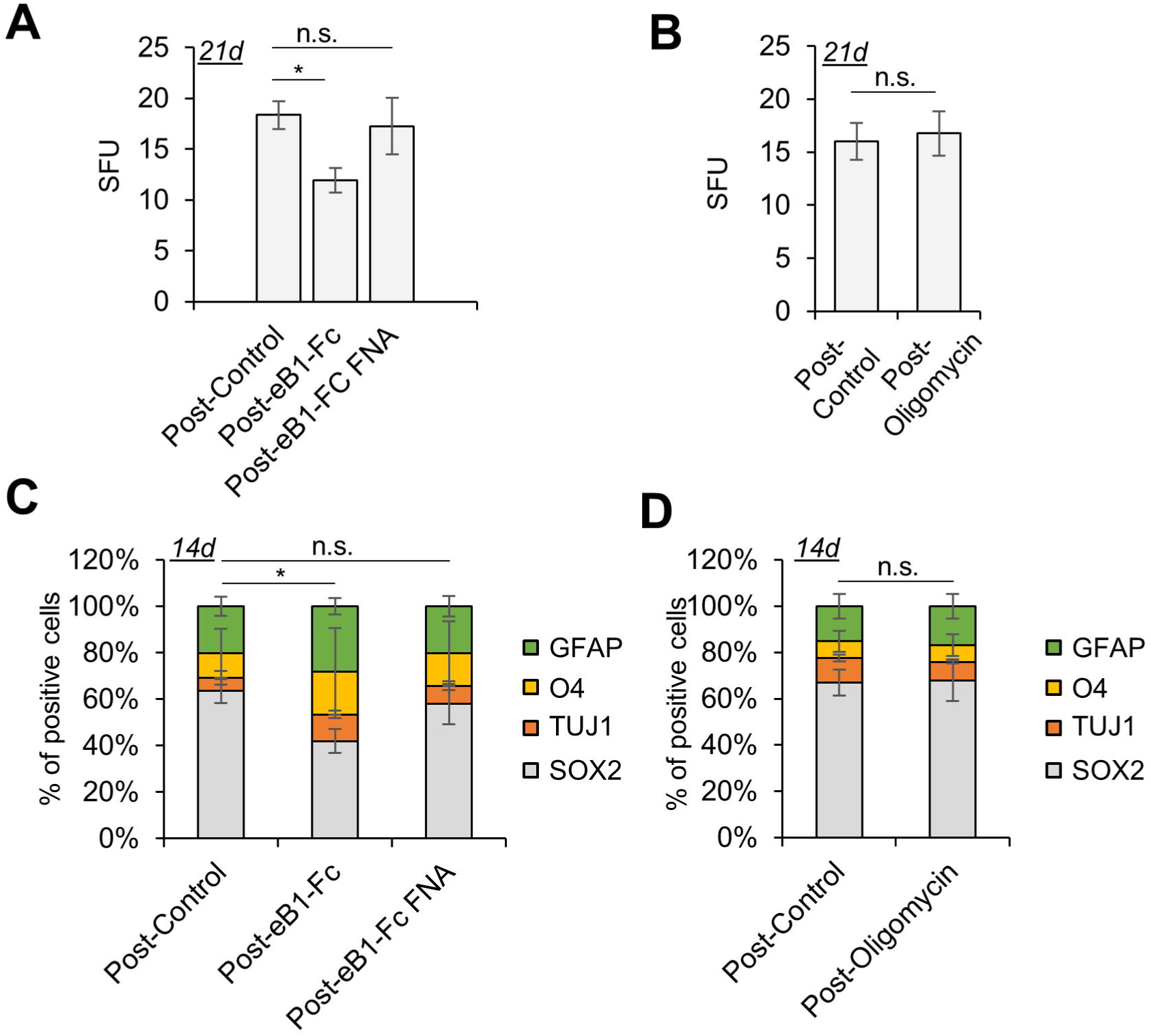
Long term inheritance of the differentiation program following DHFR inhibition. (A and B) Neurospheres were dissociated and treated with either eB1-Fc or eB1-Fc supplemented with FNA (A) or Oligomycin (B) for 7 days then dissociated and cultured under normal culture conditions for one week. Neurospheres were then dissociated and then cultured under stem culture conditions (10^2^ cells/ml) for 1 week and the number of spheres were counted. (C and D) Quantification by FACS analysis of neurospheres treated with eB1-Fc supplemented or not with FNA (C) or Oligomycin (D) for one week and cultured without treatment for another week. Cells were stained with SOX2, TUJ1, O4 and GFAP antibodies. Statistical significance was performed on SOX2 population. Statistical analysis was performed using one-way ANOVA test followed by the Bonferroni method (A), or Fisher’s LSD test (C) and Mann whitney test (B, D). Data are reported as mean ± SEM (*P < 0.05).

### Eph-B forward signaling induces NSC differentiation via the folate pathway in the developing neocortex

Next, we thought to assess whether regulation of DHFR expression by Eph-B forward signaling was relevant to NSC self-renewal vs. differentiation in vivo. First we mined a recently published single-cell RNA-sequencing (RNA-seq) dataset of mouse embryonic cortical progenitors and their progeny (Okamoto et al., 2016). These bioinformatics analyses revealed a correlation between decreased DHFR expression and neuronal differentiation (Figure S5A). This is consistent with previously published global analysis of gene expression in neural progenitors revealing that the folate pathway is upregulated in NSC compared to their differentiated progeny (Karsten et al., 2003). Furthermore, analysis of DHFR expression within the apical progenitor population revealed an inverse correlation with Eph-B2 expression. Indeed, cells with high EphB2 expression exhibited the lowest DHFR expression (Figure S5B) strengthening the link between Eph-B forward signaling and DHFR. Next, in order to confirm this link genetically, we generated Efnb1/Efnb2 double knock out embryos (dKO) and analyzed them at E13.5, a developmental stage at which the neocortex is mostly composed of neural progenitors. Western-blot analyses of neocortex tissue show that as expected, phosphorylation of Eph-B receptors is decreased in dKO embryos indicating that Eph-B signaling is turned off (Figure 6A). Importantly, this inactivation of Eph-B signaling in dKO embryos correlated with an increase in DHFR and in H3K4me3 levels compared to wild type embryos (Figure 6B). Finally, we detected an increase in both SOX2 and Nestin in dKO embryos strengthening the link between DHFR/H3K4me3 levels and the expression of these progenitor-related genes (Figure 6B). These data clearly demonstrate the molecular link between Eph-B forward signaling, DHFR expression and levels of H3K4me3 in vivo.

**Figure 6.**
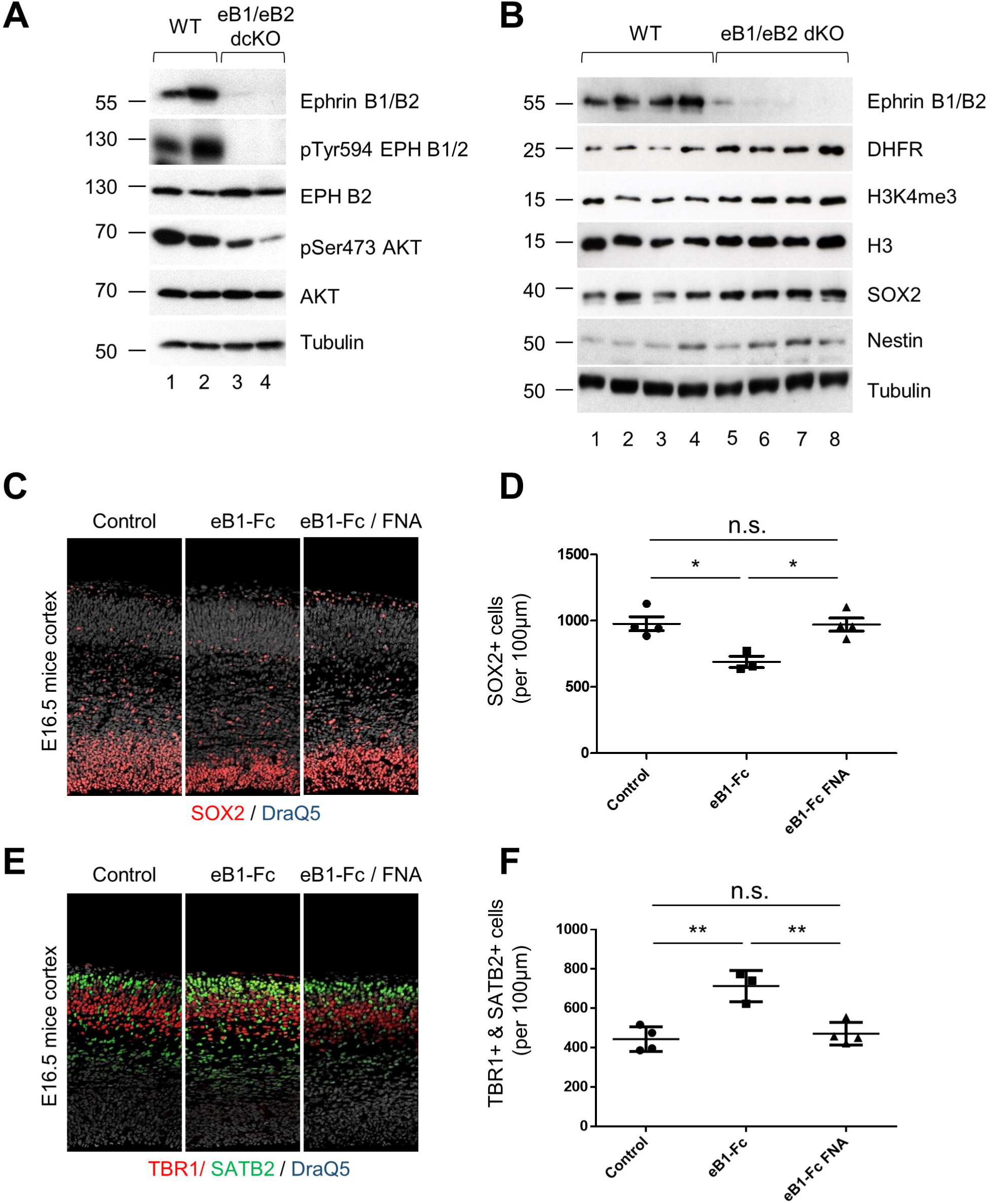
Eph B forward signaling induces NSC differentiation via the folate pathway in vivo. (A) Western blot analysis of E13.5 wild type (WT) and *Efnb1^−/−^; Efnb2^lox/lox^; nestin-Cre* (eB1/eB2 dKO) mice. Embryonic neocortex were lysed and immunoblotted with the indicated antibodies. (B) Representative confocal images of E16.5 coronal sections of embryonic neocortex 72h after in utero injection of eB1-Fc or eB1-Fc and FNA and stained for SOX2 (red) and DAPI (blue). (C) Quantification of SOX2+ cells from (B). (D) Representative confocal images of E16.5 coronal sections of embryonic neocortex 72h after in utero injection of eB1-Fc or eB1-Fc and FNA and stained for TBR1 (red), SATB2 (green) and DraQ5 (grey). (E) Quantification of TBR1+/SATB2+ cells from (B). Statistical analysis was performed using one-way ANOVA test followed by the Bonferroni method (C, E). Data are reported as mean ± SEM (*P < 0.05, **P < 0.01).

Next, to validate the role of the Eph-B forward signaling/folate axis in the control of NSC maintenance and differentiation in vivo, we injected eB1-Fc or eB1-Fc with FNA into the ventricles of E13.5 embryos in utero and analyzed the progenitor and neuronal fractions at E16.5. eB1-Fc treatment led to a decreased number of SOX2+ cells in the ventricular zone of the neocortex and this was rescued upon addition of FNA (Figure 6C and D). Furthermore, the decrease in the number of progenitor cells in response to eB1-Fc was correlated with an increase in the number of TBR1+ and SATB2+ neurons in the cortical plate (Figure 6E and F). Consistent with in vitro data, FNA supplementation rescued the effect of eB1-Fc treatment on the number of neurons (Figure 6E and F) but had no significant effect alone (Figure S5C an D). These in vivo results corroborate the in vitro data and highlight a link between 1C folate metabolism, Eph-B forward signaling and NSCs differentiation during neocortex development.

## DISCUSSION

During development, the generation of an organized tissue requires coordinated mechanisms to regulate the differentiation of cells at the appropriate time and place. Differentiation, driven by the implementation of a genetic program, correlates tightly with the execution of a metabolic program allowing for a more efficient energy production to match the evolving metabolic demands of an increasingly specialized progeny. Indeed, variations in metabolism have been reported to impact not only on cell growth and proliferation but also on lineage progression and differentiation (Sieber and Spradling, 2017). While cross talks between metabolic and signaling pathways have been well documented for cell growth and proliferation (Folmes et al., 2012a; Vander Heiden et al., 2011), how metabolic pathways are regulated to drive stem cells along specific lineages remain to be defined. Here, we show that local cell-to-cell communication regulates 1C metabolism in NSC with consequences on cell fate (Figure 7).

**Figure 7.**
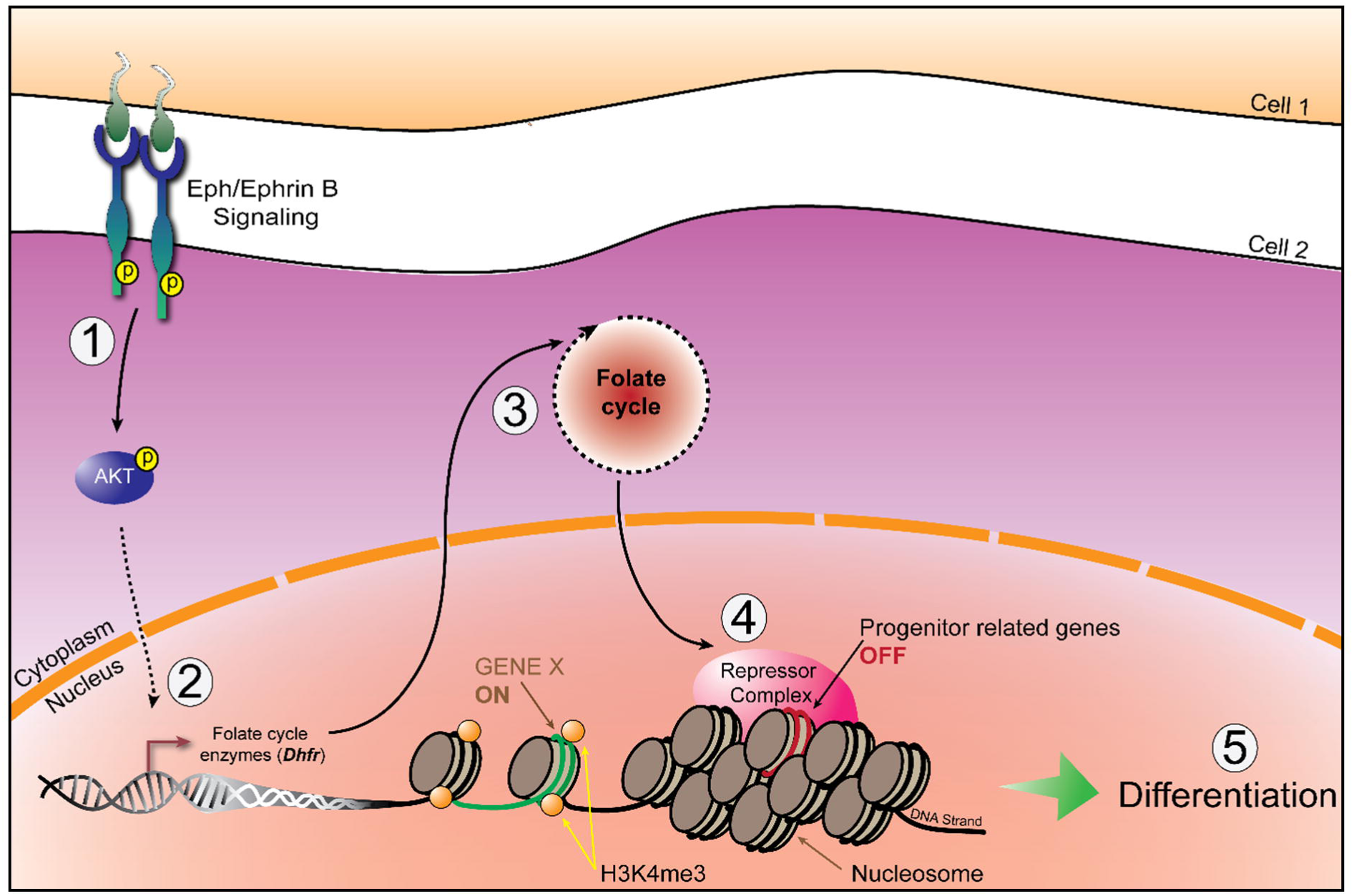
Schematic model. Schematic diagram illustrating the role of Eph-B forward signaling in the regulation of DHFR activity and the folate-dependent epigenetic program that dictates self-renewal vs differentiation of NPCs.1; Eph B forward signaling activation and phosphorylation of its downstream effector AKT, 2; reduction of DHFR expression and activity, 3; reduction of the folate pathway output, 4; decrease of H3K4me3 on the promoter of progenitor related genes, 5; decrease in the progenitor pool and increase in differentiation.

The 1C pathway plays essential roles in major metabolic processes including nucleic acid biosynthesis and methylation reactions. One of the key co-factor of the pathway is folate (also known as vitamin B12) which in animals is provided through nutrition. It has been known for many years that folate deficiency during pregnancy yields a higher risk of neural tube defect (Schorah et al., 1980). More recently, a human syndrome characterized by low folate levels in the cerebrospinal fluid (Cerebral Folate Deficiency) has been described and shown to correlate with neurological disorders. Genetic causes for this syndrome are associated with mutations in genes coding for enzymes or transporters of the 1C folate pathway (Serrano et al., 2012). For instance, mutations in *DHFR* have been reported to cause severe neurological disorders including microcephaly, the latter being consistent with an alteration in the production or survival of neurons during fetal life (Banka et al., 2011; Cario et al., 2011).

Our data show that forced activation of Eph signaling in NSC leads to down regulation of *DHFR* and increased differentiation while loss of Eph signaling correlates with increased levels of DHFR in vivo. Our results are consistent with the fact that expression of *DHFR* is downregulated as neocortex neural progenitors differentiate into neurons (Okamoto et al., 2016)Within the developing neocortex, NSC are exposed to folate through the cerebrospinal fluid and they express Eph receptors and ephrins, including ephrin-B1 (Arvanitis et al., 2013). We propose that variations in ephrin expression, either as a function of position or time, or upon cell differentiation (Arvanitis et al., 2010), modulate *DHFR* expression in neighboring cells, thus modifying epigenetic marks and contributing to prime NSC for differentiation. One difference between in vitro and in vivo analyses are the kinetics of differentiation, indeed increased differentiation was observed after 3d of eB1-Fc treatment in vivo but only 7d in vitro. This could be due to the fact that all of the in vitro analyses were performed in culture conditions that favor self-renewal whereas NSC in the developing neocortex are exposed to differentiative cues. Interestingly, our in vitro data shows that once modified, the epigenetic landscape and the pro-differentiative state of NSC is maintained in the long term, suggesting that signaling events occurring early in the neocortex developmental sequence may have consequences at later time.

The 1C pathway has two major branches, the Methionine and DNA synthesis branches that act on methylation reaction and cellular proliferation respectively. Since no apoptosis and cellular proliferation defects were observed, our results indicate that activation of Eph signaling mainly perturbs the Methionine branch of the 1C pathway in NSC. Indeed, in vitro stimulation of Eph signaling led to a decrease in H3K4me3 levels while other Histone3 methylation were not strongly affected. Such a specific decrease in H3K4me3 levels has been observed following perturbation of the Methionine cycle via Methionine or Threonine deprivation (Mentch et al., 2015; Shiraki et al., 2014; Shyh-Chang et al., 2013b). Thus it was proposed that regardless of the nutrient source, maintenance of the H3K4me3 levels in stem cells requires high SAM levels, while the threshold required for other Histone methylation appears to be lower. Interestingly, we show that decrease of H3K4me3 was only detected at promoters of progenitor-related genes and not differentiation-related genes. Metabolites can act as cofactors for epigenetic modifications and selectivity may result from gene-specific recruitment of Histone methyltransferase or Histone demethylase that have differential response to metabolites fluctuations (van der Knaap and Verrijzer, 2016).

It has been shown in mouse embryonic stem cells (mESC) that the global decrease in H3K4me3 levels following Threonine deprivation correlated with a decreased expression of pluripotency genes, including SOX2 (Shyh-Chang et al., 2013b). In fact, modulation of the Methionine cycle has been shown to impact stem cell maintenance vs. differentiation both in mESC and in human induced pluripotent stem cells (hiPS) (Shiraki et al., 2014; Shyh-Chang et al., 2013b). These studies, together with our results, indicate that maintenance of pluripotent or somatic stem cell fate requires high activity of the Methionine branch of 1C metabolism. They further suggest that stifling of 1C metabolism could be an integral part of differentiation programs. In conclusion, our data highlight a molecular cascade stretching from the cell surface to the cellular transcriptional program of NSC. Whether coupling between 1C metabolism and local cell-to-cell communication is relevant in other types of stem cells remains to be determined.

## STAR METHODS

Detailed methods are provided in the online version of this paper and include the following:

- KEY RESOURCES TABLE
- CONTACT OF REAGENT AND RESOURCE SHARING
- EXPERIMENTAL MODEL AND SUBJECT DETAILS

- Animals
- Culture of NSC and maintenance
- METHOD DETAILS

- Stimulation of Eph Forward Signaling
- Microarray Analysis
- Cell Proliferation Assay
- Sphere Forming Assay
- Quantitative PCR
- In utero injections
- Immunofluorescence and Flow Cytometry
- Protein Extraction and Western Blot
- DHFR Catalytic Assay
- Chromatin Immunoprecipitation Protocol
- Genotyping
- STATISTICAL ANALYSIS
- DATA AND SOFTWARE AVAILABILITY

- Data resources

## SUPPLEMENTAL INFORMATION

Supplemental information includes 4 figures and 1 table.

## AUTHOR CONTRIBUTIONS

M.A.F. and A.D. conceptualized the study, M.A.F. designed, performed and analyzed experiments, and co-wrote the manuscript. T.J. assisted and performed FACS analysis. T.J. performed the micro array experiment and J.S.I. analyzed the micro-array data. C.A. assisted and performed qRT-PCR experiments on NSC. C.A. performed in utero injections and A.K performed IF on mice cortices. All authors contributed to the editing of the manuscript.

## ACKNOWLEDGMENTS

We thank Arnaud Besson and members of the Pituello team for feedback and P. Belenguer’s team for sharing reagents. We acknowledge core support from the Imagery Platform of Toulouse. This work was funded by ARC (Association pour la Recherche contre le Cancer; PJA 20131200200) and ANR (Agence Nationale de la Recherche; ANR-15-CE13-0010-01). M.A. Fawal was supported by an ARC fellowship. Core funding was from the Université Paul Sabatier and the CNRS.

## STAR METHODS

### EXPERIMENTAL MODEL AND SUBJECT DETAILS

#### Animals

Wild type mice kept in a 129S4/C57Bl6J mixed background were bred in the animal facility. *Efnb1^loxlox^, Efnb2^loxlox^* and *Nestin-Cre* mouse lines have been described previously (Davy *et al*., 2004; Grunwald *et al*., 2004; Tronche *et al*., 1999). To generate compound mutants, *Efnb1^loxlox^; Efnb2^loxlox^* females were bred with *Nestin-Cre; Efnb1^Ylox^; Efnb2^loxlox^* males. Neonates were individually genotyped by PCR. Animal procedures were pre-approved by the appropriate Animal Care Committee (APAFIS#1289-2015110609133558 v5).

#### Culture of NPCs and maintenance

Cultures of primary NPCs were obtained as described previously (Chojnacki and Weiss, 2008). Briefly, embryonic day 14.5 (E14.5) cortices were dissected mechanically in phosphate buffered saline solution (PBS) (Cat#D1408, Sigma). The single-cell suspension was collected, rinsed with DMEM/F12 (Cat#1130-032, Invitrogen), and cultured with growing medium (DMEM/F12 medium containing 0.6% glucose (Cat#UG3050, Euromedex), 5 mM HEPES (Cat#H3375, Sigma), 1 mM putrescine (Cat#P5780, Sigma), 5 ng/ml basic fibroblast growth factor 2 [FGF-2] ( Cat#F029, Sigma), 20 ng/ml epidermal growth factor [EGF] (Cat#E9644, Sigma), 10 ng/ml insulin-transferrin-sodium selenite supplement (Cat#I1844-1VL, Sigma) and 2% B27 supplement (Cat#17504-044,Invitrogen)) in a 5% CO2 incubator at 37°C. Several different primary cultures were obtained and kept in culture for no more than 5 passages. Fresh medium was added every 2–3 days to the culture and passage was realized once a week using Accutase^®^ (Cat#A6964, Sigma).

### METHODS DETAILS

#### Stimulation of Eph Forward Signaling

Prior to stimulation, neurospheres were dissociated with Accutase^®^ (Cat#A6964, Sigma), and aliquots of 10^5^ cells were placed in growing medium then stimulated with 1 μg/ml eB1-Fc (Cat#473-EB, R&S Systems) preclustered with 0.1 μg/ml anti-human IgG (Cat#G-101-C-ABS, R&D Systems). Total RNA was isolated 6h, 72h post-stimulation and processed for qRT-PCR. Alternatively, cells were stimulated for 30min, 6h, 24h and 72h and lysed in protein lysis buffer for Western blot analysis.

#### Microarray Analysis

MouseWG-6_V2_0_R3_11278593_A.txt was obtained from Illumina and used to extract Illumina identifiers (column: Array_Address_Id) and probe sequences (column: Probe_Sequence), which were converted to nuIDs using lumi:seq2id() (Du et al., 2007, 2008). The limu:lumiMouseAllACCNUM R object was used to map nuIDs to RefSeq accessions. By connecting through the nuID, we were then able to assign RefSeq accessions and relevant gene identity information to each probe on the microarray. The readIDAT_enc() function from illumiaio (Smith et al., 2013) was used in order to read the Grn.idat binary files and obtain the TrimmedMeanBinData, which was assembled into a matrix for downstream limma analyses (Ritchie et al., 2015). Quantile normalization was applied with limma:normalizeBetweenArrays and limma was used to optain log2 Fold Change and p-values for each contrast (lmFit, contrasts.fit and eBayes). To identify genes differentially expressed upon forward signaling activation, the forward-2hour, forward-6hour and control-6hour data were contrasted against the control-2hour time (n = 3 for each group). The differential expression between control-6hour and control-2hour was used to evaluate the time-dependent changes in gene expression, independently of treatment. P-values were adjusted via limma:topTable (Benjamini & Hochberg method) and all results were assembled into data.frame for further queries by biologists using a spreadsheet program.

#### Cell Proliferation Assay

Neurospheres were dissociated with Accutase^®^ (Cat#A6964, Sigma), and cells were seeded in triplicate in a 6-well plate at 10^4^ cell/ml density. Cell number was measured using Cellometer™ (Cat#1001201, Nexcelom) according to the manufacturer’s instructions.

#### Sphere Forming Assays

Neurospheres were dissociated with Accutase^®^ (Cat#A6964, Sigma), and about hundred cells were seeded in a 96-well plate, and the total number of spheres was counted 2 weeks after incubation. eB1-Fc (Cat#473-EB, R&S Systems), Methotrexate (Cat#M8407, Sigma) and Oligomycin (Cat#O4876, Sigma) were added to the assay culture medium upon seeding to a final concentration of 1 μg/ml, 2 μM and 0.1 μg/ml respectively. Sphere-Forming Unit (SFU) was calculated: SFU = (number of spheres counted / number of input cells) * 100.

#### Quantitative PCR

RNA was extracted from cell pellets using TRIreagent (Cat#TR118, MRC) according to the manufacturer’s instructions. 1 µg RNA was used for reverse transcription. Genomic DNA was degraded with 1 µl DNase (Cat#M6101, Promega) for 20 min at 37°C in 20 µl RNase/DNase-free water (Cat#W4502, Bio-RAD), and the reaction was stopped by adding 1 µl stop solution under heat inactivation at 65°C for 10 min. 2 µl dNTPs (10 mM; Promega) and 2 µl oligodTs (100 mM, idtDNA) were added for 5 min at 65°C, then 8 µl of 5× buffer, 2 µl RNasin (Cat#N2511, Roche), and 4 µl of 100 mM DTT (Cat#P1171, Promega) were added for 2 min at 42°C. The mix was divided into equal volumes in a reverse-transcriptase–negative control tube with addition of 1 µl water and in a reverse-transcriptase–positive tube with 1 µl superscript enzyme and placed at 42°C for 1 h. The reaction was stopped at 70°C for 15 min, and cDNAs were diluted (10-, 100-, and 1,000-fold) and processed for quantitative PCR in triplicate for each dilution. 10 µl diluted cDNA was mixed with 10 µl premix Evagreen (Cat# 1725204, Bio-RAD) containing 1µM of each primer, and the PCR program was run for 40 cycles on a CFX96 BioRad Realtime system (Cat#185-5096, Bio-RAD). mRNA relative expression levels were calculated using the 2-ΔΔCts method. Primers are listed in Table 1.

#### In utero injections

Timed-pregnant mice were anesthetized using Vetflurane (Cat#Vnr137317, Vibrac) and uterine horns were exposed. DMSO, 1 µg/ml MTX (Cat#M8407, Sigma), 1μg/ml pre-clustered human IgG-Fc (Cat#G-101-C-ABS, R&D Systems) or 1µg/ml pre-clustered eB1-Fc (Cat#473-EB, R&S Systems) were injected in the lateral ventricle of E13.5 embryos neocortex using pulled micropipettes. Body wall cavity and skin were sutured and embryos were allowed to develop normally for 72 h. Embryonic brains were collected at E16.5, fixed with 4% paraformaldehyde (Cat#15710, Fisher scientific) overnight at 4°C. Vibratome sections (50 µm) (Cat#VT1000 S, Leica Biosystems) were processed for immunofluorescence staining as described below.

#### Immunofluorescence and Flow Cytometry

For immunofluorescence, cells were grown and treated as described earlier on. Neurospheres were collected and fixed at room temperature with 4% paraformaldehyde (Cat#15710, Fisher Scientific) for 30 min, then permeabilized and blocked with PBTA solution (0.5% BSA (Cat#04100811C, Euromedex), 0.5% FBS (Cat#10500064, ThermoFisher), and 0.1% Triton X-100 (Cat#T9284, Sigma) in PBS (Cat#D1408, Sigma) for 30 min. Primary and secondary antibodies diluted in PBTA were incubated overnight at 4°C and 2h at room temperature, respectively. Neurospheres were collected with centrifugation at 1,200 rpm for 10 min at each step. Neurospheres were finally mounted on glass slides (Superfrost) (Cat#LR90SF03, ThermoFisher) with Coverslips using mounting medium (4.8% wt/vol Mowiol (Cat#81381, Sigma) and 12% wt/vol glycerol (Cat#G9012, Sigma) in 50 mM Tris pH 8.5 (Cat#26-128-3094-B, Euromedex)).

Cortices were permeabilized and blocked with PBTA2 solution (2% BSA, 2% FBS, 1% Tween20 (Cat#2001-A, Euromedex) for 2h at room temperature. Primary antibodies diluted in PBTA2 and secondary antibodies diluted in PBS were incubated about 36h at 4°C and 1.5h at room temperature respectively. Labelled sections were finally mounted on glass slides (Superfrost) with Coverslips using mounting medium (4.8% wt/vol Mowiol and 12% wt/vol glycerol in 50 mM Tris, pH 8.5).

Microscope acquisitions were performed on either an SP5 DM600B confocal microscope (40×/1.3 APO; 20×/0.7 APO, PMT; Leica Biosystems) or an SP8 droit (40x, TCS; Leica Biosystems). Observation was performed using Type-F mineral oil (1153859; Leica Biosystems). Cell counting and pixel quantification were performed with the use of ImageJ software.

For fluorescence-activated cell sorter (FACS) analysis, neurospheres were dissociated with Accutase^®^ (Cat#A6964, Sigma) and approximately 10^6^ cells in 1.2ml ice-cold PBS were fixed in 4% PFA. The cell suspension was incubated in 200µl PBTA first with either anti-Tuj1 and anti-SOX2 antibodies or anti-GFAP and anti-O4 antibodies and then with a secondary antibody coupled to AlexaFluor 488 and 647 for overnight at 4°C and 2h at room temperature respectively. Cells were resuspended in 500µl PBS and acquired with a FACSCalibur cytometer (Cat#342975, Becton Dickinson). To set the threshold for specific positive fluorescence intensity signal, we use control samples with cells incubated with secondary antibodies alone.

#### Protein Extraction and Western Blot

For protein extraction, Neurospheres were pelleted by centrifugation at 1,500 rpm for 10 min and washed twice with an ice-cold PBS (Cat#D1408, Sigma) solution. Whole cell extracts were obtained by re-suspending cell pellets in the following ice-cold lysis buffer (150 mM NaCl (Cat#1112-A, Euromedex), 50 mM Tris-HCl pH 7.4 (Cat#EU0011, Euromedex), 0.5 mM EDTA (Cat#EU0007, Euromedex), 2 mM Na3VO4(Cat#S6508, Sigma), 1% Nonidet P-40 (NP-40) (Cat#N6507, Sigma), 0.5 mM EGTA (Cat#E4378, Sigma) and 0.1 mM PMSF (Cat#78830, Sigma)) supplemented by protease inhibitors (Cat#**11836170001, Roche)** for 1h on ice. Protein lysates were then vortexed, sonicated with Biorupter^®^ Sonicator for 10 cycles (Cat#B01020001, Diagenode), and centrifuged at 13,000 rpm for 10 min. Cleared lysates were either used for Western blot analyses. Samples were denatured by boiling in loading buffer (4× 100 mM Tris-HCL, pH 6.8, 8% SDS (Cat#EU0660, Euromedex), 40% glycerol (Cat#G9012, Sigma), 4% β-mercaptoethanol (Cat#63689, Sigma), and bromophenol blue (Cat#B0126, Sigma)) before loading and electrophoresis on an 8 or 16% SDS-PAGE gel. Proteins were transferred onto a nitrocellulose membrane (Cat#**GE10600002,** GE Healthcare), which was blocked for 30 min and incubated with primary antibody in 5% nonfat dry milk in TBS-T (20 mM Tris base (Cat#26-128-3094-B, Euromedex), 150 mM NaCl, and 0.05% Tween 20 (Cat#2001-A, Euromedex) adjusted to pH 7.6 with 1 M HCl (Cat#30721, Sigma)) overnight at 4°C. Milk was replaced with 5% BSA (Cat#04100811C, Euromedex) for detection of phosphorylated epitopes. Western blots presented are representative of at least three independent experiments.

#### DHFR Catalytic Assay

DHFR activity, measured as folate reductase, was determined by the dihydrofolate reductase assay kit according to the manufacturer’s instructions. Briefly, cells were pelleted by centrifugation at 1,500 rpm for 10 min and washed twice with an ice-cold PBS solution (Cat#D1408, Sigma). Cell pellets were resuspended in 300µl assay buffer supplemented with protease inhibitors (Cat#**11836170001, Roche)** and lysed mechanically using glass beads (Cat#G8772, Sigma). Lysates were precleared by centrifugation at 13,000 rpm for 10 min. The supernatant was used immediately for enzyme assay after the determination of protein concentration.

#### Chromatin Immunoprecipitation Protocol

NPCs were treated as described above and cultured for 72h. Cells were fixed with 1% formaldehyde (Cat#15710, Fisher Scientific) for 15 min at room temperature, with occasional swirling. Glycine (Cat#G7126, Sigma) was added to a final concentration of 0.125 M and the incubation was continued for an additional 5 min. Cells were collected and washed with ice-cold PBS (Cat#D1408, Sigma) three times and resuspended in cell lysis buffer (5 mM Pipes [pH 8] (Cat#P6757, Sigma), 85 mM KCl (Cat#P017-A, Euromedex), 0.5% NP-40 (Cat#N6507, Sigma), Protease inhibitors (Cat**#11836170001, Roche)**. Cells were centrifuged at 4000 rpm for 10 min at 4°C. Pellets were then resuspended in Nuclei lysis buffer (50 mM Tris-HCl [pH 8] (Cat#EU0011, Euromedex), 10 mM EDTA, 1% SDS). The cells were disrupted by sonication (30 cycles, 30 secs on, 60 secs off) with Biorupter^®^ Sonicator (Cat#B01020001, Diagenode). The chromatin solution was clarified by centrifugation at 15,000 g at 4°C for 10 min. The average DNA fragment size was 250 pb. The chromatin solution was diluted with IP dilution buffer (16.7 mM Tris-HCl [pH 8], 1.2 mM EDTA (Cat#EU0007, Euromedex), 1.1 mM Triton X-100 (Cat#T9284, Sigma), 0.01% SDS (Cat#EU0660, Euromedex) and 167 mM NaCl (Cat#1112-A, Euromedex) and pre-cleared with pre-blocked beads (protein G Sepharose (Cat#3296, Sigma)/protein A agarose (Cat#6526, BioVision) (50/50) overnight with PBS/0.5% BSA (Cat#04100811C, Euromedex), 10mg/ml yeast tRNA (Cat#AM7119, ThermoFisher) beads for 1h at 4°C. The pre-cleared diluted chromatin sample was incubated with 3 μg of anti-H3K4me3 (Cat#ab8580, Abcam) overnight at 4°C. Preblocked beads were added for an additional 4h. The beads were washed twice with the dialysis buffer (2 mM EDTA, 50 mM Tris-HCl [pH 8] and 0.2% N-lauroylsarcosine (Cat#L5777, Sigma), 4 times with wash buffer (100 mM Tris-HCl [pH 8], 500 mM LiCl (Cat#L4408, Sigma), 1% NP-40 and 1% sodium Deoxycholate (Cat#D6750, Sigma) and twice with TE buffer (10 mM Tris-HCl [pH 8] and 1 mM EDTA). The immunoprecipitated material was eluted from the beads by heating for 15 min at 65°C in 1% SDS, 50 mM NaHCO3 (Cat#71631, Sigma). 0.2 M NaCl (Cat#1112-A, Euromedex) and 10μg/ml RNAse A (Cat#EN0531, ThermoFisher) were added before incubating for 2h at 64°C to reverse the crosslinks. Samples were then incubated with 1.5 μg/ml Proteinase K (Cat#P6556, Sigma), 40 mM Tris-HCl [pH 8], 10 mM EDTA at 45°C for 1h. The samples were then extracted with phenol chloroform isoamyl alcohol (Cat#0038.2, Carl ROTH) followed by ethanol (Cat#20821.330, VWR chemicals) precipitation in the presence of glycogen (Cat#R0561, ThermoFisher), and resuspended in DNAse-free water (Cat#W4502, Sigma). The resulting precipitated DNA was amplified and analyzed by qRT-PCR as described above.

### STATISTICAL ANALYSIS

For experiments involving a single pair of conditions, statistical significance between the two sets of data were analyzed with a Mann-Whitney and Wilcoxon test with Prism5 (GraphPad software). For datasets containing more than two samples, one-way analysis of variance with either Bonferroni or Fisher’s LSD multiple comparison post-test was used to determine adjusted p-values. Sample sizes of sufficient power were chosen on the basis of similar published research and were confirmed statistically by appropriate tests. Each experiment was performed at least three times, and 100–500 cells were counted for each condition of each experiment. Statistically significant differences are reported at P < 0.05, P < 0.01, P < 0.001.

### DATA AND SOFTWARE AVAILABILITY

#### Data resources

The accession number for the microarray data reported in this paper is GEO: GSE104068.

**Table 1:**
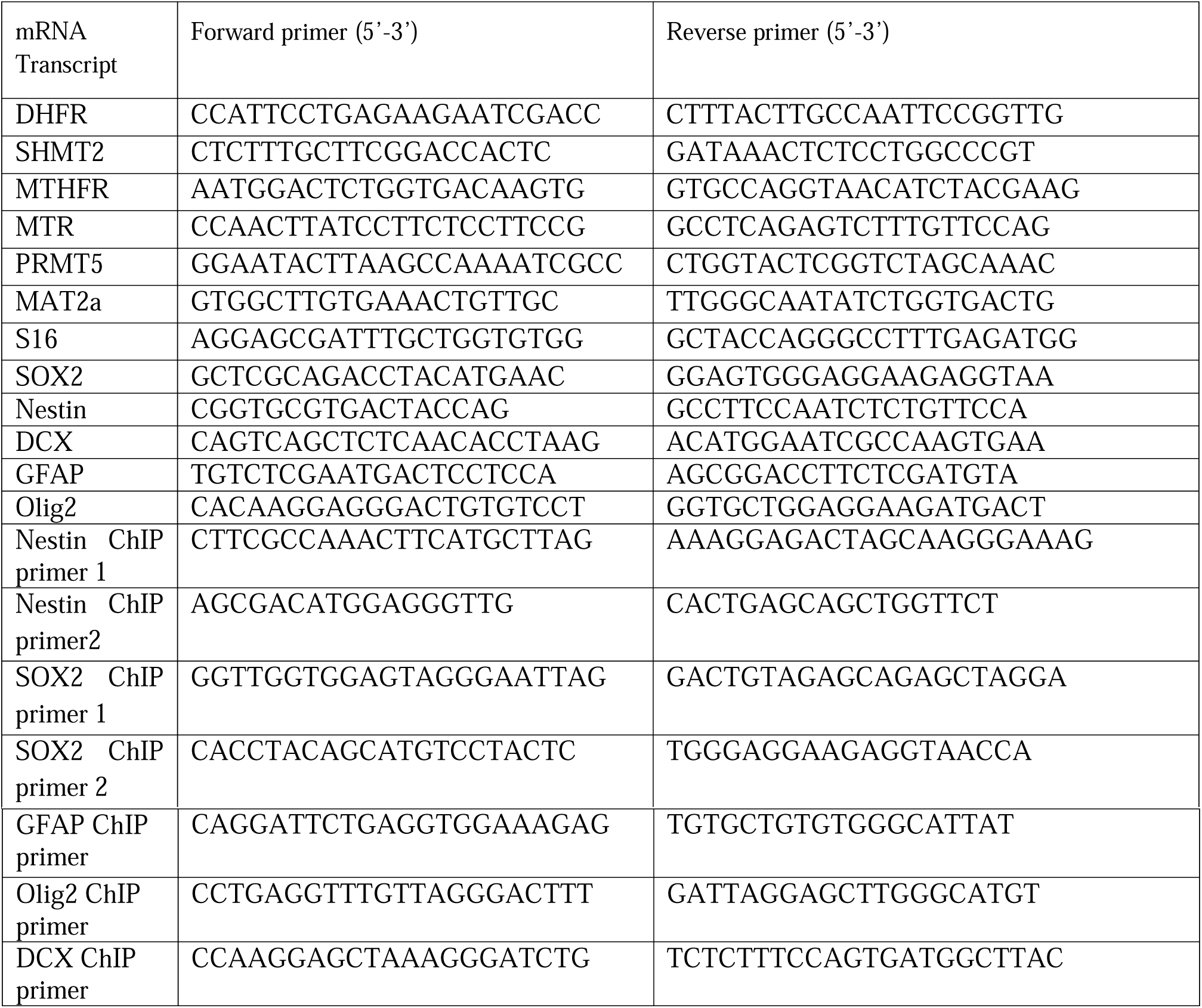
Primer sequence used for quantitative RT-PCR analysis. Related to STAR Methods.

